# Competing valence-related roles of dopamine in the tail of the striatum

**DOI:** 10.1101/2025.08.06.668813

**Authors:** Ryota Tsuruga, Yu Tajika, Masabumi Minami, Iku Tsutsui-Kimura

## Abstract

Understanding how the brain evaluates aversion and appetition to guide behavior remains an open question. Here, we investigated the role of dopamine signaling in the tail of the striatum (TS) in regulating competing valence-based behaviors and learning. TS dopamine dynamics were monitored as mice performed a classical conditioning task in which an odor cue predicted either an aversive air puff or a water reward. Initially, mice exhibited anticipatory blinking, which diminished over time, while anticipatory licking emerged later, coinciding with adaptation to the air puff. Dopamine responses in the TS to the air puff and its associated odor were initially elevated but declined with repeated exposure. Optogenetic disruption of this decline suppressed adaptation and hindered appetitive learning. These findings demonstrate that TS dopamine dynamics are essential for behavioral adaptation to aversive stimuli, which indirectly facilitates appetitive learning, underscoring a regulatory mechanism for shifting between defensive and reward-seeking behaviors.

## Introduction

Animals must often make decisions to obtain rewards while avoiding aversive or threatening stimuli, such as foraging within a predator’s territory. Appetitive behavior and learning depend on environmental context. In the absence of threat, individuals can engage in diverse behavioral patterns and explore across multiple dimensions. This increases the likelihood of encountering rewards and encoding appetitive information along with relevant environmental cues. In contrast, in threatening environments, attention shifts toward aversive stimuli, reducing motivation for reward-seeking and disrupting the acquisition of appetitive learning. Although substantial research has examined how aversive stimuli interfere with appetitive learning and behavior^1–4^, the underlying brain mechanisms remain largely unknown.

Dopamine dynamics within the mesolimbic pathway play a central role in appetitive learning and approach behavior^5,6^. Recent studies have identified a distinct subpopulation of dopamine neurons projecting to the tail of the striatum (TS), which receive unique presynaptic inputs^7^ and exhibit specialized activity. These neurons respond to novel objects, multiple sensory modalities, and threat-associated cues^8–12^, but do not consistently signal reward in rodents and monkeys^10,13^. Ablation of dopamine signals in the TS reduces avoidance of aversive or threatening stimuli^8,11^, suggesting that TS dopamine contributes to avoidance of negative events. Despite limited reward-related signaling, some studies have shown that TS dopamine encodes reward values^13,14^ or modulates performance in reward-driven choice tasks^15^, resembling canonical dopamine function. Therefore, clarifying the precise roles of TS dopamine in processing aversive and appetitive stimuli is critical for understanding the neural basis of competing valences.

In this study, we monitored and manipulated dopamine activity in the TS during aversive and appetitive learning in mice. To this end, we developed a two-valence classical conditioning (2-VCC) task in which an odor served as a conditioned stimulus (CS), followed by either a water reward (USwater) or an aversive air puff (USpuff) as the unconditioned stimulus (US). We found that TS dopamine responded to both the US and CS during air puff trials (CSpuff), with responses gradually declining across trials. Because this decline correlated with adaptation to the USpuff, we optogenetically prevented the decay of the USpuff response to examine behavioral effects. Stimulation inhibited adaptation to the aversive stimulus and disrupted appetitive learning during the adaptation phase. These findings indicate that dopamine signaling in the TS is essential for adapting to aversive events and, in turn, supports appetitive learning.

## Results

### Two-valence classical conditioning task for studying the interaction of aversive and appetitive learning

To investigate how interference between two competing valences affects learning and behavior, we developed a two-valence classical conditioning (2-VCC) task in which water-deprived mice were trained to associate a neutral odor CS with either an aversive air-puff or a water reward (Fig. 1A–B). The task included three trial types: appetitive, aversive, and null (Fig. 1C). These were presented in a pseudo-randomized order with a fixed ratio of 3:1:3. Mice underwent 70 trials per session, one session per day, over 5 consecutive days. Blinking and licking behaviors were recorded as indices of aversive and appetitive responses, respectively.

**Fig.1.**
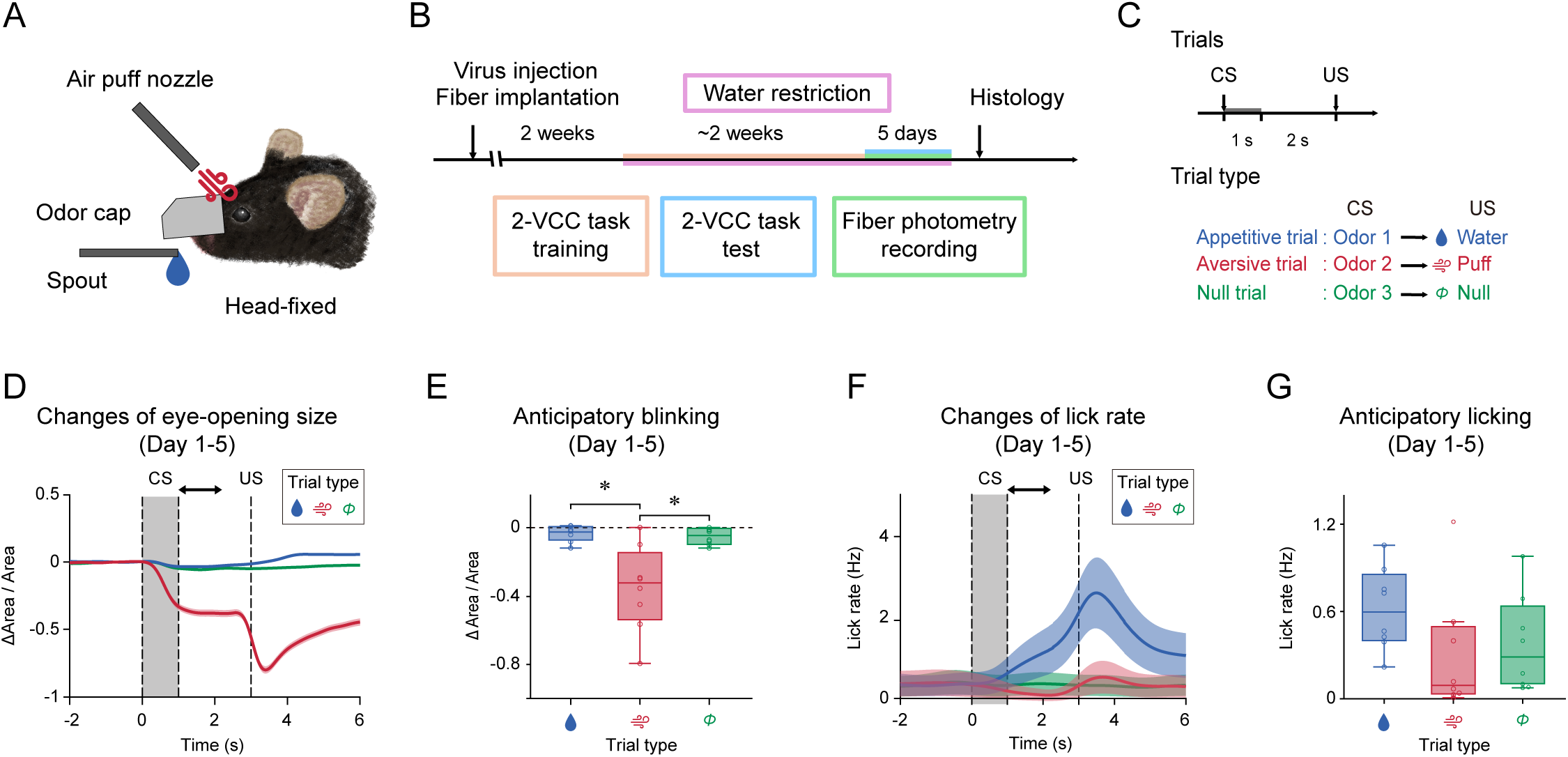
2-VCC task evaluates aversive and appetitive learning simultaneously. **A,** Schematic of the 2-VCC task. **B,** Schedule of the behavioral experiment. **C,** Time-course of the 2-VCC task. **D,** Peristimulus time histogram (PSTH) of eye-opening size (average of 5 sessions). The arrow indicates the analysis time-window (1 s – 2 s from the CS presentation). **E,** Mice discriminatively showed anticipatory blinking in the aversive trial (right, p=1.6×10^-4^, F(2, 14)=17.3, repeated ANOVA; p=4.0×10^-3^, aversive vs appetitive, p=5.0×10^-3^, aversive vs null, p=0.065, appetitive vs null, Bonferroni’s test, n=8 animals for each). **F,** PSTH of lick rate (average of 5 sessions). The arrow indicates the analysis time-window (1 s – 2 s from the CS presentation). **G,** There was no significant difference in anticipatory licking across trial types (right, p=0.018, F(2, 14)=5.4, repeated ANOVA; p=0.054, aversive vs appetitive, p=1.0, aversive vs null, p=0.13, appetitive vs null, Bonferroni’s test, n=8 animals for each). Error bars, SEM. Center of box plot shows median; edges are 25th and 75th percentiles; and whiskers are the most extreme data points. *p < 0.05.

We concatenated the 5-days dataset (Fig. 1D–G and S1) and confirmed that mice blinked in response to air-puff stimuli and licked during water delivery (Fig. 1D and F, Video S1–2). Notably, mice exhibited blinking prior to the US only in aversive trials (Fig. 1E, Video S3), indicating anticipatory blinking as a prospective defensive response to the impending aversive stimulus^16,17^.

Similarly, an increased anticipatory lick rate was observed during appetitive trials (Fig. 1G). This behavior became more pronounced in later sessions (days 3–5 and Fig. S1), suggesting that mice developed anticipatory licking as a prospective approach behavior toward the expected appetitive stimulus^16,18^. These findings indicate that mice successfully acquired both aversive and appetitive learning in the 2-VCC task.

### The level of aversion correlates with the intensity of appetitive behavior

To examine time-dependent dynamics of aversive learning, we analyzed blinking behavior across sessions and trials (Fig. 2A–B). Contrary to our initial hypothesis that aversive learning would intensify over time, anticipatory blinking weakened, particularly in later sessions (Fig. 2B). Based on these patterns, we divided the sessions into two phases (sessions 1–2 vs. 3–5) and assessed how anticipatory defensive behavior evolved across trials within each phase (Fig. 2C–D). During the early phase, anticipatory blinking increased across trials, suggesting that mice exhibited avoidance learning by actively preparing for the aversive stimulus (Fig. 2C). In the later phase, anticipatory blinking diminished across trials, indicating adaptation learning, in which mice acclimated to repeated exposure to the aversive stimulus (Fig. 2D). Therefore, we designated sessions 1–2 as the avoidance phase and sessions 3–5 as the adaptation phase of aversive learning.

**Fig.2.**
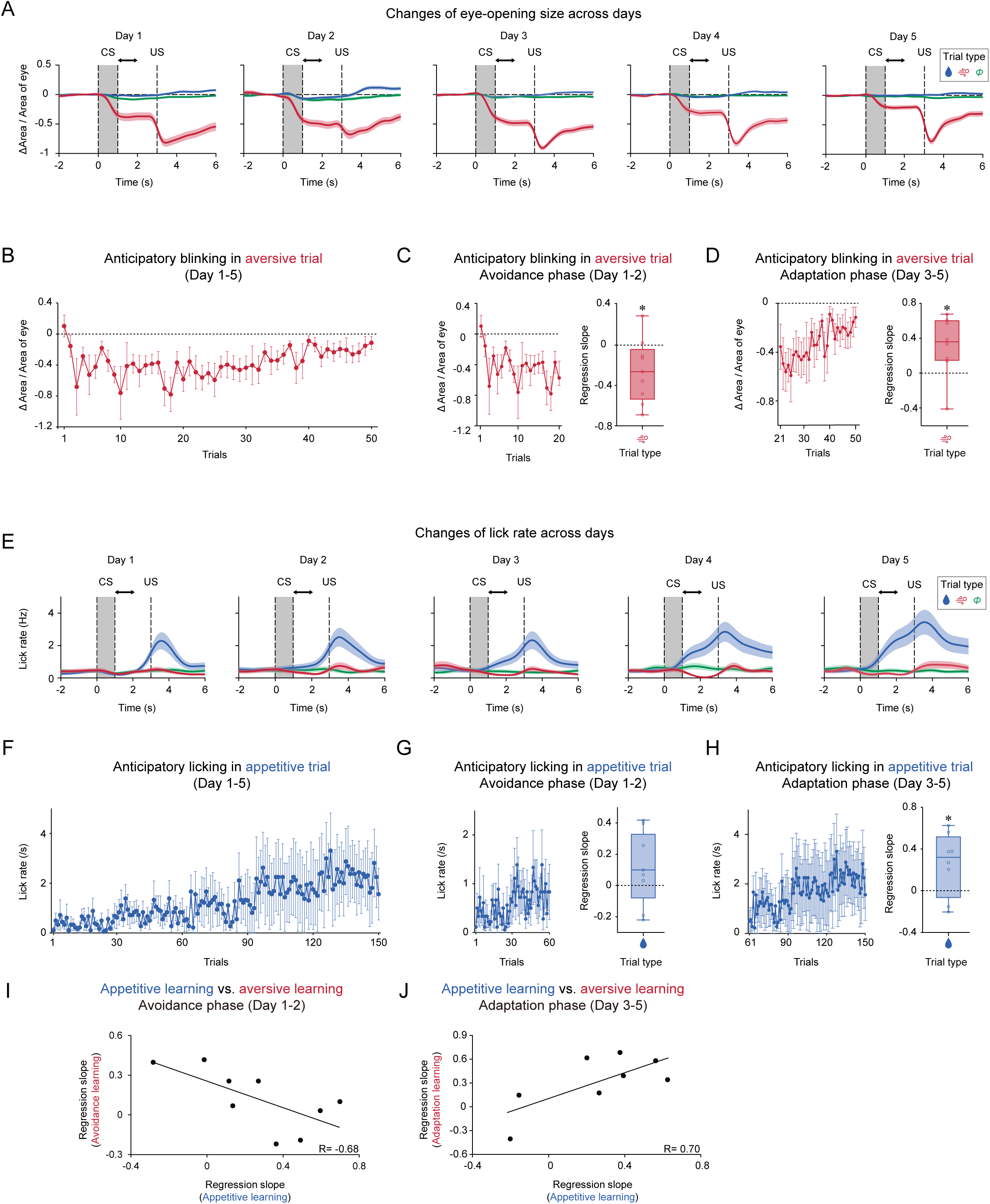
Time-dependent changes of anticipatory blinking and licking. **A,** PSTHs of eye-opening size across days. The arrow indicates the analysis time-window (1 s – 2 s from the CS presentation). **B,** Time-course of anticipatory blinking in the aversive trials. **C,** Regression coefficients of anticipatory blinking with trial number in Day 1-2 for each animal were significantly negative (p=0.035, one-sampled t-test, n=9). **D,** Regression coefficients of anticipatory blinking with trial number in Day 3-5 for each animal were significantly positive (p=0.039, one-sampled t-test, n=8). **E,** PSTHs of lick rate across days. The arrow indicates the analysis time-window (1 s – 2 s from the CS presentation). **F,** Time-course of anticipatory licking in the appetitive trials. **G,** No significant change in regression coefficients of the rate of anticipatory licking with trial number in Day 1-2 for each animal (p=0.16, one-sampled t-test, n=9). **H,** Regression coefficients of the rate of anticipatory licking with trial number in Day 3-5 for each animal are significantly positive (p=0.047, one-sampled t-test, n=8). **I,** Regression coefficients of levels of avoidance learning with appetitive learning were anti-correlated (R=-0.68, p=0.044, Pearson’s correlation coefficient, n=9). **J,** Regression coefficients of levels of adaptation learning with appetitive learning were correlated (R=0.70, p=0.052, Pearson’s correlation coefficient, n=8). Error bars, SEM. Center of box plot shows median; edges are 25th and 75th percentiles; and whiskers are the most extreme data points. *p < 0.05.

We then examined the time-dependent dynamics of appetitive learning by analyzing licking behavior across sessions and trials (Fig. 2E–F). Although anticipatory licking increased across trials (Fig. 2E), no statistically significant increase was observed across sessions 1–5 (Fig. 2F and S2A). To explore the relationship between aversive and appetitive learning, we applied the aversive learning phase classification to appetitive learning analysis. Anticipatory licking remained unchanged during the avoidance phase (Fig. 2G) but significantly increased during the adaptation phase (Fig. 2H), indicating a potential interaction between the two learning modalities. We further examined correlations between aversive and appetitive learning intensities in both phases and found that their regression slopes were negatively correlated during the avoidance phase but positively correlated during the adaptation phase (Fig. 2I–J and S2B). These results suggest that appetitive learning was suppressed when mice developed defensive behavior in anticipation of aversive stimuli but enhanced when they adapted to repeated aversive exposure. In summary, aversive learning can be stratified into two phases—avoidance and adaptation—and the intensity of appetitive learning negatively correlates with avoidance and positively correlates with adaptation.

### TS dopamine discriminates aversive conditioned stimulus

To determine whether dopamine signaling in the striatum influences aversive and appetitive learning, we recorded dopamine dynamics in the tail of the striatum (TS) and the anterior striatum (AS) by injecting an AAV virus expressing the GRAB_DA_ sensor into the respective subregions (Fig. 3A, D, and S3A). Multi-bundle fiber photometry was used to measure dopamine signals in both the TS and AS of the same animal. Recordings were conducted while mice performed the 2-VCC task over 5 days (Fig. 1B). Dopamine signals were concatenated across the 5 recording days (Fig. 3B and E). We observed that average TS dopamine levels increased in response to CSpuff and the aversive stimulus itself (Fig. 3B), whereas AS dopamine showed no clear response (Fig. 3E). Quantitative analysis revealed a significant increase in TS dopamine release—but not in the AS—selectively during aversive trials (Fig. 3C and F). We then assessed the relationship between recording site locations and the magnitude of dopamine CS responses (Fig. 3G and S3B), finding a clear correlation between anterior–posterior positioning and TS dopamine response to CSpuff. These findings indicate that TS dopamine has a unique role in encoding aversive information, consistent with previous reports^9,10^.

**Fig.3.**
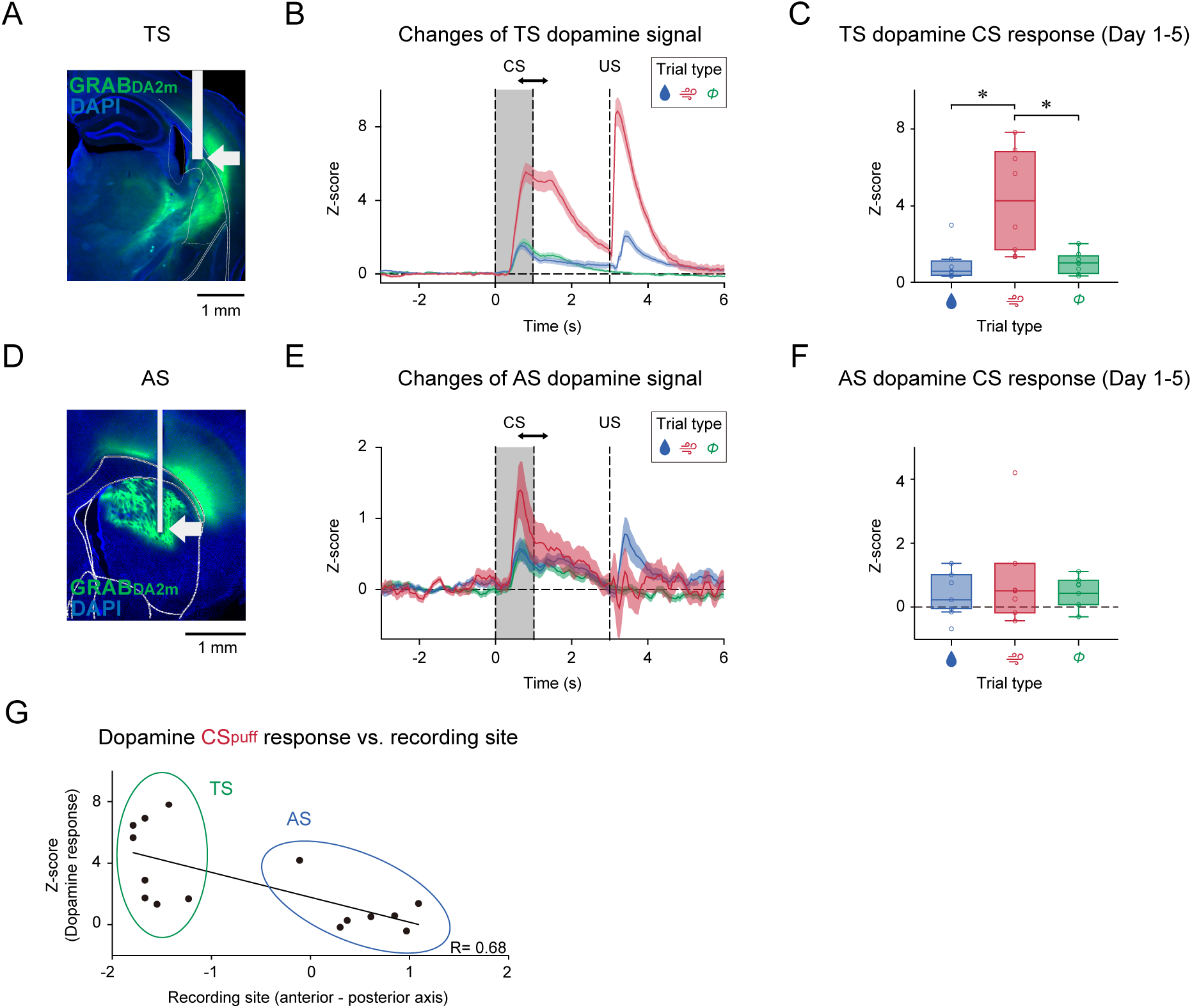
TS dopamine discriminates CSpuff. **A,** AAV9-hSyn-GRAB_DA2m_ was injected unilaterally into TS of WT mice. Representative image of GRAB_DA2m_ expression in the TS. White arrow, the tip of a fiber. Green,GRAB_DA2m_. Blue, DAPI. Bar, 1 mm. **B,** PSTH of dopamine response (average of 5 sessions). The arrow indicates the analysis time-window (0.5 s – 1.5 s from the CS presentation). **C,** CS response of TS dopamine was discriminatively enhanced in the aversive trial (p=2.0×10^-4^, F(2, 14)=16.71, one-way ANOVA; p=0.013, aversive vs appetitive, p=0.013, aversive vs null, Bonferroni’s test, n=8 animals for each). **D,** AAV9-Syn-GRAB_DA2m_ was injected unilaterally into AS of WT mice. Representative image of GRAB_DA2m_ expression in the AS. White arrow, the tip of a fiber. Green,GRAB_DA2m_. Blue, DAPI. Bar, 1 mm. **E,** PSTH of dopamine response (average of 5 sessions). The arrow indicates the analysis time-window (0.5 s – 1.5 s from the CS presentation). **F,** CS response of AS dopamine was equivalent across trial types (p=0.45, F(2, 12)=0.85, one-way ANOVA, n=7 animals for each). **G,** Dopamine CS response was higher in the more posterior striatum (R=-0.68, p=5.0×10^-3^, Pearson’s correlation coefficients, n=15 animals). Error bars, SEM. Center of box plot shows median; edges are 25th and 75th percentiles; and whiskers are the most extreme data points. *p < 0.05.

### TS dopamine gradually decays during adaptation to aversion

We next investigated how dopamine responses evolved across different phases of aversive learning (Fig. 4A–D). Initially, TS dopamine responses to CSpuff were high but declined gradually over trials (Fig. 4A–B). We then examined TS dopamine dynamics during the previously defined avoidance and adaptation phases (Fig. 4C–D). During the avoidance phase, TS dopamine responses to CSpuff remained elevated across trials (Fig. 4C). By contrast, a significant decrease in TS dopamine response was observed during the adaptation phase (Fig. 4D).

**Fig.4.**
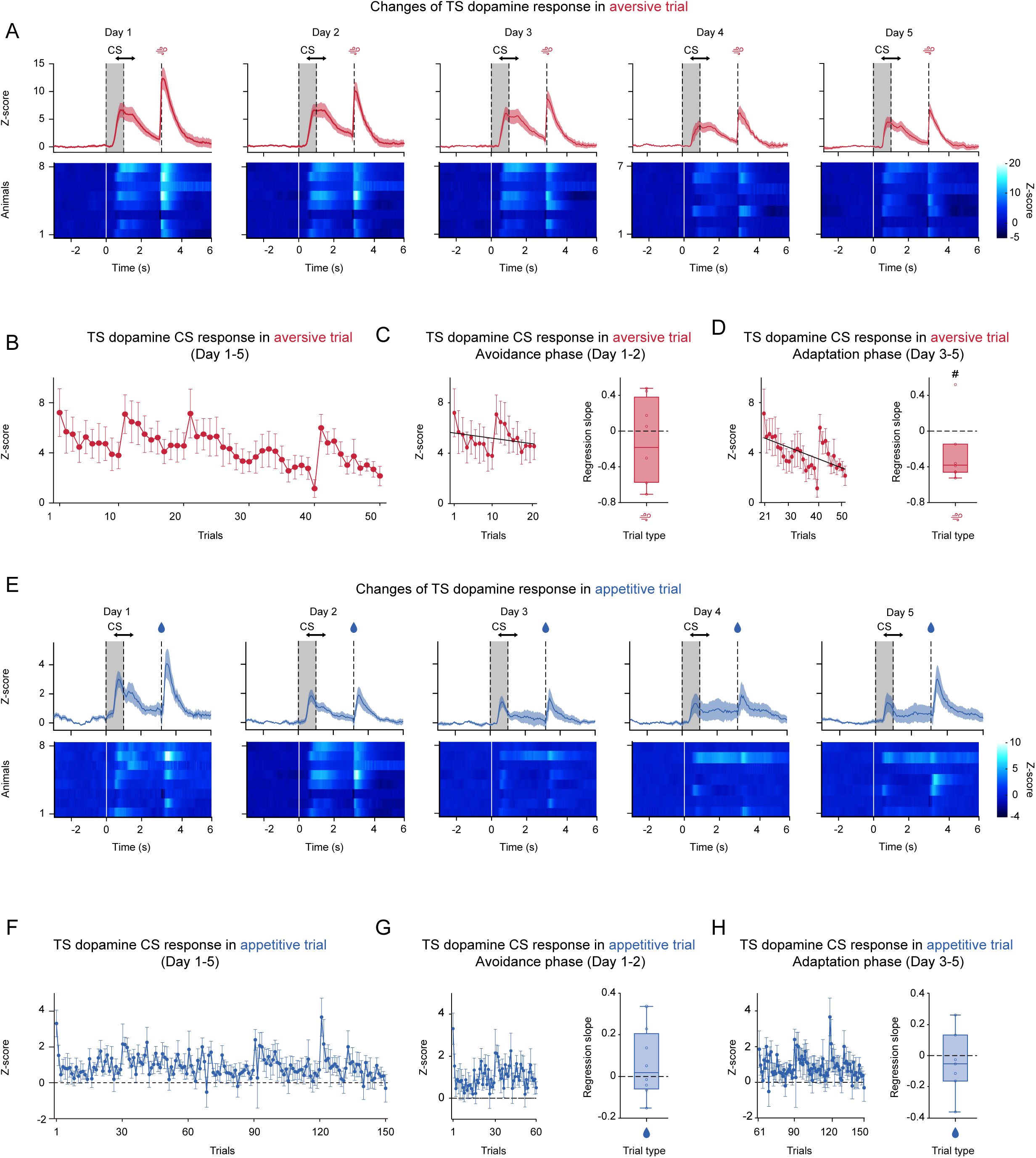
TS dopamine dynamics in the aversive and appetitive trials. **A,** PSTHs and heat maps of TS dopamine response in the aversive trials across days. The arrow indicates the analysis time-window (0.5 s – 1.5 s from the CS presentation). **B,** Time-course of CSpuff response of TS dopamine in the aversive trials. **C,** No significant change in regression coefficients of CSpuff response with trial number in Day 1-2 for each animal (p=0.47, one-sampled t-test, n=8). **D,** Regression coefficients of CSpuff response with trial number in Day 3-5 for each animal are significantly positive (p=5.0×10^-3^, one-sampled t-test, n=7). **E,** PSTHs and heat maps of TS dopamine response in the appetitive trials across days. The arrow indicates the analysis time-window (0.5 s – 1.5 s from the CS presentation). **F,** Time-course of CSwater response of TS dopamine in the appetitive trials. **G,** No significant change in regression coefficients of CSwater response with trial number in Day 1-2 for each animal (p=0.56, one-sampled t-test, n=8). **H,** No significant change in regression coefficients of CSwater response with trial number in Day 3-5 for each animal (p=0.34, one-sampled t-test, n=7). Error bars, SEM. Center of box plot shows median; edges are 25th and 75th percentiles; and whiskers are the most extreme data points. *p < 0.05.

We also analyzed TS dopamine dynamics during appetitive learning (Fig. 4E– H). TS dopamine showed a modest increase in response to CSwater and USwater (Fig. 4E), in agreement with a previous report^9^. Unlike responses to CSpuff, TS dopamine signals to CSwater did not exhibit dynamic changes (Fig. 4F), regardless of whether trials occurred in the avoidance (Fig. 4G) or adaptation phase (Fig. 4H).

These findings suggest that although TS dopamine does not encode appetitive information, its decline during the adaptation phase may contribute to acclimatization to repeated aversive stimuli.

### Adaptation development correlates with TS dopamine decay

Given the observed decrease in TS dopamine signaling over trials during the adaptation phase (Fig. 4D), we hypothesized that this decline contributes to adaptation to aversive stimuli. To test this, we analyzed correlations between the regression slopes of behavioral adaptation and TS dopamine dynamics (Fig. 5A–B and S4). Mice exhibiting greater decreases in TS dopamine responses tended to show stronger adaptation (Fig. 5B). Supporting this, anticipatory blinking was reduced in trials with lower dopamine signaling during adaptation (Fig. S4). No clear correlation was found between the degree of avoidance learning and TS dopamine changes (Fig. 5A). Additionally, no significant correlation was observed between appetitive learning intensity and changes in TS dopamine signaling during either learning phase (Fig. 5C–D). These results suggest a potential role for TS dopamine signaling in the development of adaptation to aversive stimuli.

**Fig.5.**
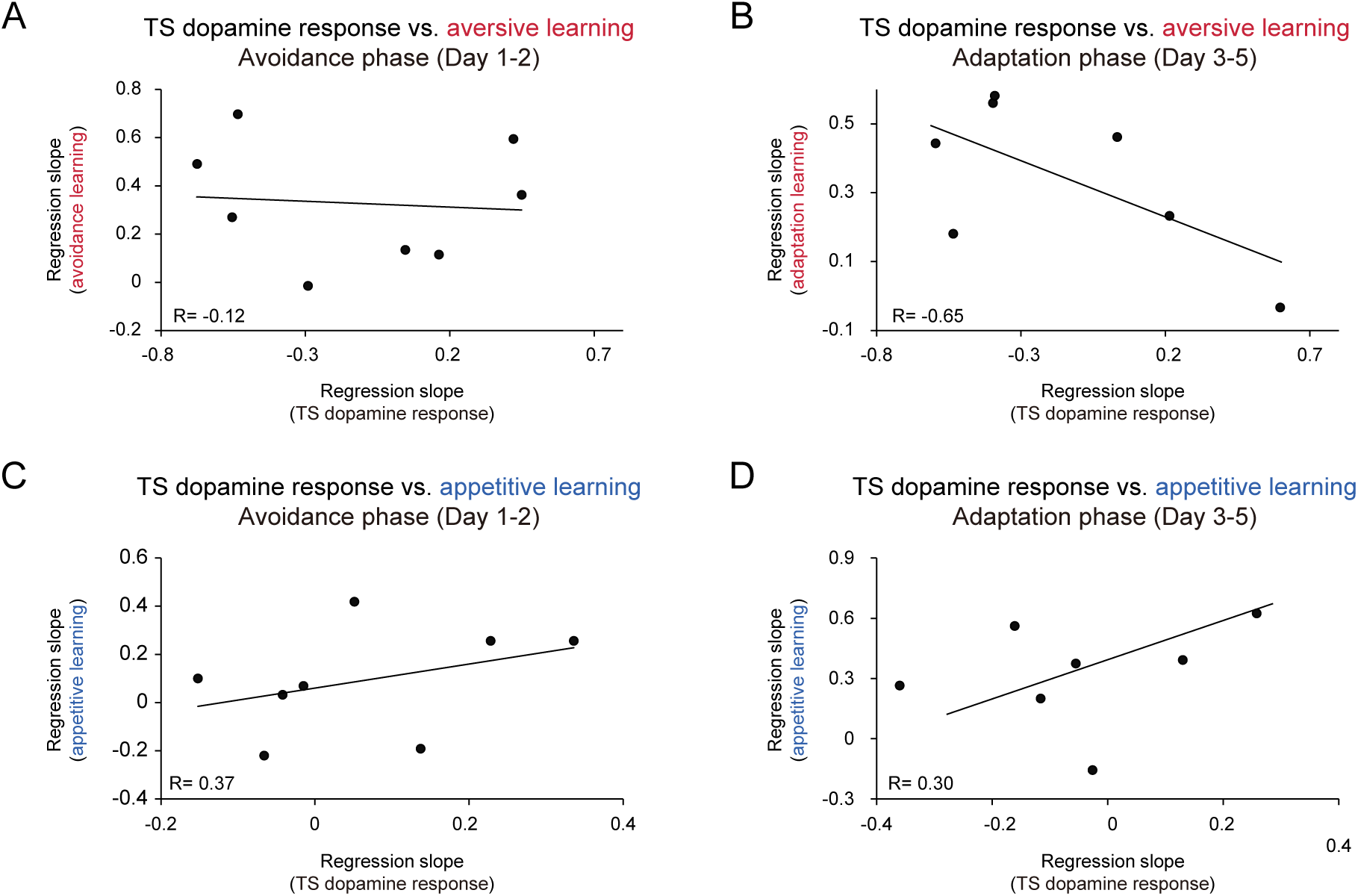
Correlations between TS dopamine changes and performance of learning. **A,** Regression coefficients of levels of avoidance learning with changes of TS dopamine CSpuff response were not correlated (R=-0.12, p=0.78, Pearson’s correlation coefficient, n=8). **B,** Regression coefficients of levels of adaptation learning with changes of TS dopamine CSpuff response in the avoidance phase were anti-correlated (R=-0.65, p=0.12, Pearson’s correlation coefficient, n=7). **C,** Regression coefficients of levels of appetitive learning with changes of TS dopamine CSwater response were not correlated (R=0.37, p=0.37, Pearson’s correlation coefficient, n=7). **D,** Regression coefficients of levels of appetitive learning with changes of TS dopamine CSwater response in the adaptation phase were not correlated (R=0.30, p=0.51, Pearson’s correlation coefficient, n=7).

### Decay of TS dopamine promotes adaptation to aversion and approach to reward

We previously observed that greater development of appetitive learning (Fig. 2J and S2B) and more pronounced decay of TS dopamine signaling (Fig. 5B and S4) occurred in mice exhibiting greater adaptation to aversive events. Based on these findings, we hypothesized that the decay of TS dopamine facilitates adaptation to aversion and indirectly enhances appetitive learning, despite TS dopamine not directly encoding appetitive information (Fig. 3C, 4E–H, and 5C–D). To test this hypothesis, we optogenetically manipulated TS dopamine release to counteract its decline during the adaptation phase and evaluated the resulting effects on both aversive adaptation and appetitive learning.

We targeted the dopamine response to USpuff rather than to CSpuff, given that, according to reinforcement learning theory, the US response is a driver of learning, while the CS response is considered a consequence of learning^6,19–21^. First, we optimized the frequency of optical stimulation to replicate the physiological dopamine response to air puff stimulation. Head-fixed mice expressing ChrimsonR or tdTomato in the unilateral SNL—where TS-projecting dopamine neurons reside—also expressed GRAB_DA2m_ in the ipsilateral TS (Fig. 6A and S5A–C). Mice were exposed to air-puff stimulation or red-light pulses at 5, 10, or 20 Hz. We monitored TS dopamine dynamics during the task (Fig. 6B– C). The dopamine response evoked by 10 Hz light stimulation most closely resembled the physiological response to the air puff (Fig. 6B). Notably, 10 Hz stimulation did not induce artifacts in control mice (Fig. 6B). Quantitative analysis confirmed that dopamine response to 10 Hz light was equivalent to the response to air-puff stimulation, while tdTomato controls showed no response (Fig. 6C). Based on these findings, we selected 10 Hz red-light pulses to manipulate the USpuff response.

**Fig.6.**
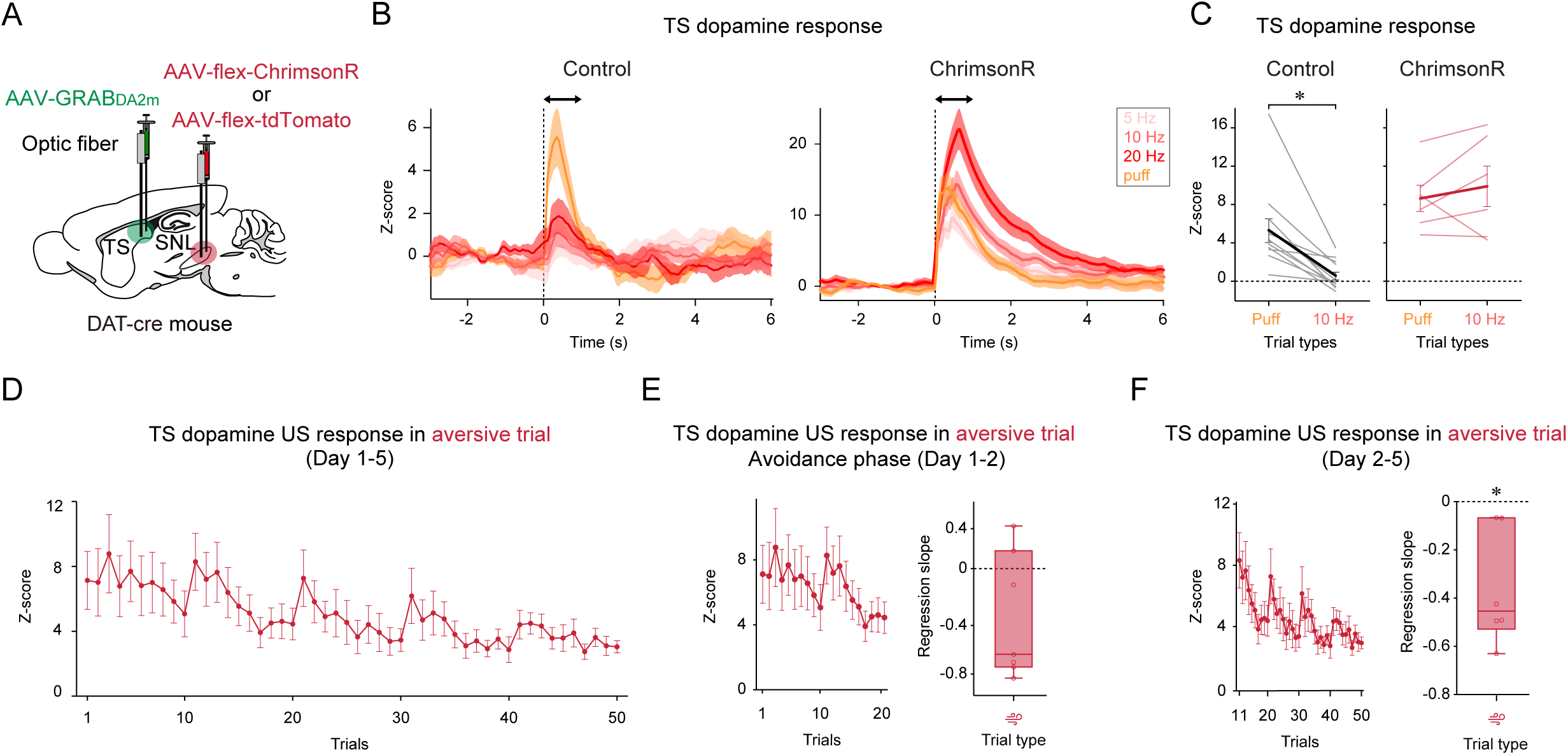
Optimization of optogenetic manipulation to mimic physiological dopamine response to air-puff. **A,** AAV9-hSyn-GRABDA2m was injected unilaterally into TS and AAV9-CAG-flex-ChrimsonR or AAV5-CAG-flex tdTomato was injected into the ipsilateral SNL of dopamine trans porter (DAT)-cre mice. **B,** PSTHs of TS dopamine response to optogenetic stimulation or air-puff application in the control (left) and ChrimsonR (right) groups. **C,** 10Hz of optogenetic stimulation yielded TS dopamine response equivalent to its response to air-puff (left, p=6.9×10^-4^, two-sided t-test, control group, n=12; right, p=0.42, two-sided t-test, ChrimsonR group, n=6). **D**, Time-course of USpuff dopamine response in the aversive trials. **E**, No change in regression coefficients of USpuff response with trial number in Day 1-2 for each animal (p=0.14, one-sampled t-test, n=7). **F**, Regression coefficients of USpuff dopamine response with trial number in Day 2-5 for each animal were significantly negative (p=0.015, one-sampled t-test, n=6). Error bars, SEM. Center of box plot shows median; edges are 25th and 75th percentiles; and whiskers are the most extreme data points. *p < 0.05.

To identify the appropriate phase for manipulation, we examined TS dopamine responses to air-puff application during aversive trials across sessions 1–5. Dopamine responses to the US remained elevated during the avoidance phase and significantly declined in sessions 2–5 (Fig. 6D–F); this pattern was not evident when data from session 1 were included (Fig. S5D). Therefore, we conducted optogenetic stimulation of SNL dopamine neurons during sessions 2–5 (Fig. 7A).

**Fig.7.**
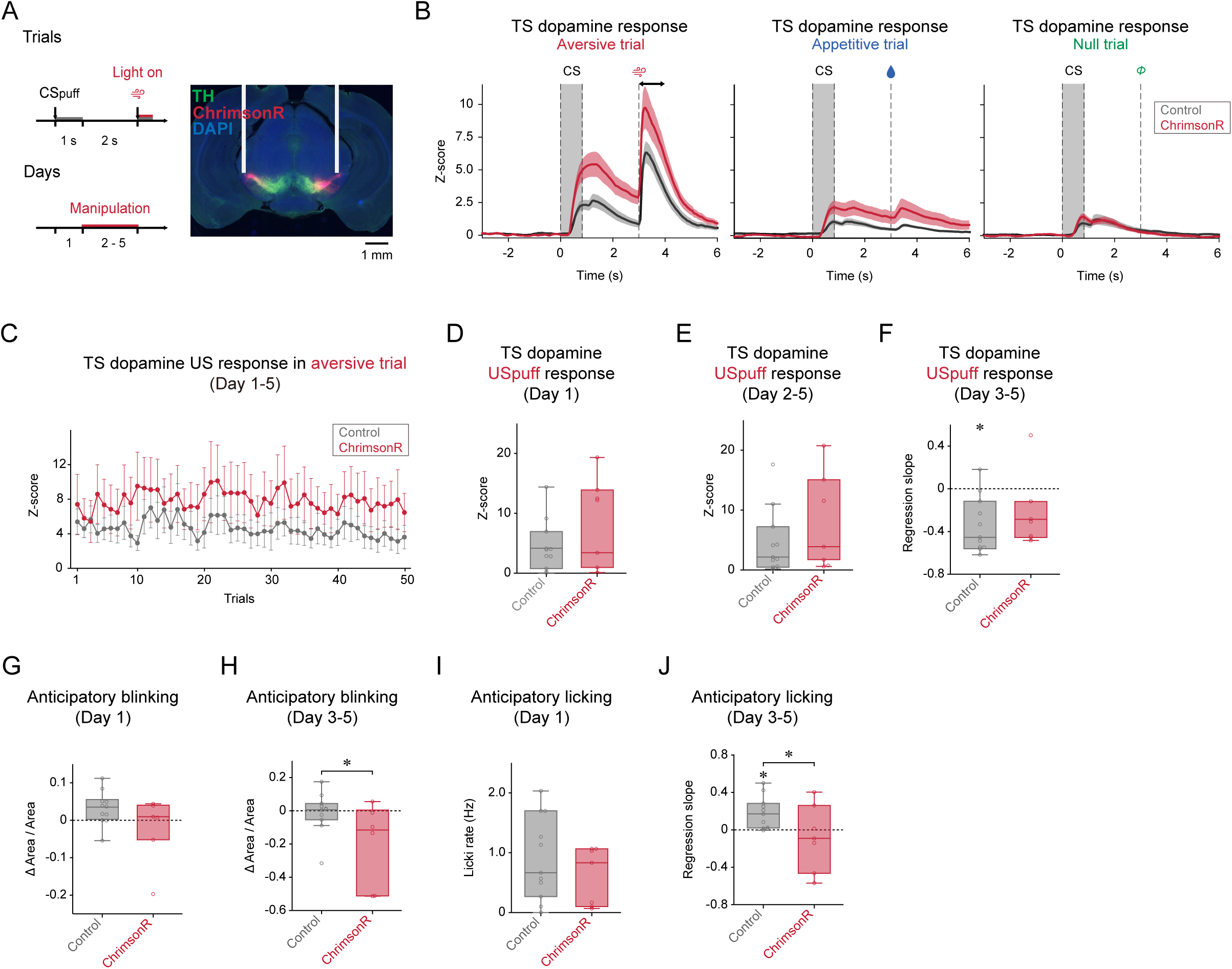
Optogenetic cancelation of TS dopamine decay affected adaptation and appetitive learning. **a**, Schedule of the optogenetic experiment and the representative image showing ChrimsonR expression in the SNL. AAV9-hSyn-GRABDA2m was injected unilaterally into TS and AAV9-CAG-flex-ChrimsonR was injected into the bilateral SNL of DAT-cre mice. Green, tyrosine hydroxylase (TH). Red, ChrimsonR. Blue, DAPI. Bar, 1 mm. **B**, PSTH of TS dopamine response in each trial types. The arrows indicate the analysis time-window (3.0 s – 4.0 s from the US presentation). **C**, Time-course of USpuff dopamine response in the aversive trials. **D**, No difference in USpuff response in control and ChrimsonR groups in Day 1 (p=0.34, two-sided t-test, control group, n=11, ChrimsonR group, n=7). **E**, USpuff response was not changed by optogenetic stimulation (p=0.30, two-sided t-test, control group, n=11, ChrimsonR group, n=7). **F**, Regression coefficients of USpuff dopamine response with trial number in Day 3-5 for each animal were significantly negative in the control (p=1.6×10^-3^, one-sampled t-test, n=11) but not in ChrimsonR group (p=0.16, one-sampled t-test, n=7). **G**, No difference in anticipatory blinking in control and ChrimsonR groups in Day 1 (p=0.14, two-sided t-test, control group, n=11, ChrimsonR group, n=6). **H**, Anticipatory blinking was significantly increased in the ChrimsonR group (p=0.048, two-sided t-test, control group, n=11, ChrimsonR group, n=6). **I**, No difference in anticipatory licking in control and ChrimsonR groups in Day 1 (p=0.36, two-sided t-test, control group, n=11, ChrimsonR group, n=6). **J**, Anticipatory licking was developed in the control ((p=8.9×10^-3^, one-sampled t-test, n=11) but not in ChrimsonR group (p=0.56, one-sampled t-test, n=7; p=0.049, two-sided t-test, control group, n=11, ChrimsonR group, n=7). Error bars, SEM. Center of box plot shows median; edges are 25th and 75th percentiles; and whiskers are the most extreme data points. *p < 0.05.

We next conducted the 2-VCC task with TS dopamine manipulation during USpuff application in sessions 2–5 (Fig. 7A and S6A-B). TS dopamine signals were combined across 5 recording days (Fig. 7B). Although optogenetic activation did not significantly enhance the USpuff response (Fig. 7C–D), it prevented the decline in TS dopamine response to USpuff during the adaptation phase (Fig. 7D–F and S6C). We then assessed the effects of optogenetic manipulation on TS dopamine response to CSpuff and found a moderate increase of TS dopamine response (Fig. S6D–G). These findings indicate that optogenetic activation during USpuff application prevented the decay of TS dopamine response to USpuff and modestly increased the CSpuff response during adaptation.

To investigate the effects of optogenetic manipulation on aversive learning, we compared the amplitude of defensive-like anticipatory blinking between intervention and control groups during aversive trials (Fig. 7G–H). On the first day, with no optogenetic manipulation, anticipatory blinking did not differ significantly between the groups (Fig. 7G). However, during the adaptation phase, the intervention group exhibited significantly greater blinking amplitude than the control group (Fig. 7H), indicating reduced adaptation to repeated aversive stimuli. These results suggest that the sustained TS dopamine response to USpuff impaired adaptation learning.

To assess the effects on appetitive learning, we analyzed anticipatory licking in appetitive trials (Fig. 7I–J). On the first day, there was no significant difference between groups in licking magnitude in the absence of optogenetic manipulation (Fig. 7I). During the adaptation phase, however, the intervention group showed a significantly lower regression slope of anticipatory licking than the control group (Fig. 7J), suggesting that impaired TS dopamine decay to CSpuff weakened appetitive learning. Together, these results suggest that TS dopamine dynamics are critical for the development of aversive adaptation and, consequently, the progression of appetitive learning.

## Discussion

Reward-related learning and appetitive behavior are influenced by the presence or absence of negative events, such as aversive or threatening stimuli. To investigate the neural mechanisms underlying interference between two opposing valences, we established a 2-VCC task designed to simultaneously evaluate aversive and appetitive learning. This was achieved by measuring blinking and licking responses as behavioral indicators of avoidance and approach, respectively. Defensive-like anticipatory blinking was frequently observed during early sessions but gradually declined in later sessions. This progression allowed us to classify aversive learning into avoidance and adaptation phases. Appetitive-like anticipatory licking emerged only during the adaptation phase, when the animals acclimated to repeated exposure to the same aversive stimuli. TS dopamine responded to USpuff and CSpuff, with these responses diminishing across trials in the adaptation phase. We counteracted this decay using optogenetic stimulation of TS-projecting dopamine neurons, which inhibited both adaptation and appetitive learning. These findings suggest that TS dopamine signaling plays a key role in accelerating habituation to aversive or threatening stimuli, thereby facilitating the development of appetitive learning.

### TS dopamine in aversive learning and adaptation

Although several studies have suggested that TS dopamine signaling contributes to associating painful stimulation with a neutral cue or to developing defensive behaviors against potential threats^8,12,22,23^, we did not observe a clear correlation between TS dopamine activity and avoidance learning in this study (Fig. 4C and 5A). This may be due to a strong initial response to CSpuff in the first trial, potentially attributable to novelty (Fig. 4B). Previous studies implemented habituation to the CS prior to conditioning, which reduced novelty-related TS dopamine responses to the CS^12,23^. Further optogenetic manipulation during the avoidance phase is necessary to elucidate the role of TS dopamine in avoidance learning within the 2-VCC task.

Consistent with previous reports ^8–10,12,22,23^, we observed that TS dopamine responded to aversive or threatening stimuli and to cues predicting such events, with responses diminishing following repeated exposure in a stable context. While most previous work has examined the role of TS dopamine in associative threat learning, Zafiri et al. (2025)^12^ assessed the effects of TS dopamine ablation on fear recall and extinction using an auditory fear conditioning task. They reported that mice with dopamine ablation could recall and extinguish fear memories, suggesting that TS dopamine is not required once the memory is formed. Interestingly, our optogenetic findings revealed that TS dopamine dynamics are essential for adaptation to aversive or threatening stimuli (Fig. 7H), a process that occurs after memory formation. Although threat extinction and adaptation share features—such as reducing fear or defensive behavior in response to a CS—they are theoretically distinct. Extinction results from procedures aimed at decreasing the threat value of the CS^24–26^ while adaptation refers to reduced threat elicited by the US^27,28^. Our result and the previous report^12^ highlight mechanistic differences between threat extinction and adaptation. Specifically, TS dopamine appears necessary for adaptation or habituation, but not for extinction.

In this study, we demonstrated the importance of TS dopamine signal decay in aversive adaptation; however, the mechanisms driving this decay remain unclear. Dopamine neurons in the lateral midbrain display greater burst activity in vivo^29^ and higher excitability in vitro^30,31^ than those in the medial midbrain, such as the ventral tegmental area (VTA), where canonical reward-processing dopamine neurons reside. These findings suggest that intrinsic or local inhibitory mechanisms may have limited involvement in TS dopamine signal reduction. Instead, the decay may arise from inhibitory afferents located outside the midbrain. Indeed, TS-projecting SNL dopamine neurons receive distinct inputs compared to other dopamine subpopulations^7^. Notably, they are innervated by the globus pallidus external (GPe) and the dorsal raphe nucleus, both of which contain GABAergic projection neurons. Future studies should investigate how dynamic dopamine signaling is regulated during the adaptation process.

### TS dopamine in appetitive learning and behavior

While optogenetic stimulation of dopamine neurons projecting to the TS inhibited appetitive learning in the present study (Fig. 7J), several prior studies involving TS dopamine ablation reported no changes in reward-driven behaviors^10,12,22^. We initially considered the possibility of unintended optogenetic activation of adjacent substantia nigra compacta (SNc) dopamine neurons. However, given that SNc dopamine neurons facilitate rather than inhibit appetitive learning^32–35^, any such stimulation would likely not account for the observed suppression. Alternatively, long-term dopamine ablation might allow for compensatory adaptations^36^, whereas our precisely timed optogenetic intervention may have more effectively revealed the role of TS dopamine in appetitive behavior. In this study, we carefully selected stimulation parameters to remain within physiological response ranges (e.g., responses to air puff stimulation, Fig. 6B–C), as recent findings indicate that physiological dopamine effects differ significantly from those induced by intense optogenetic activation^37,38^. Thus, it is also possible that unphysiological manipulations such as ablation obscured TS dopamine function in appetitive behavior.

### Potential neural circuits underlying the modulation of avoidance, adaptation, and reward acquisition

Based on current findings, TS dopamine does not appear to directly modulate appetitive learning (Fig. 4E–H and 5C–D). Instead, appetitive learning is likely governed by other brain networks, such as dopamine projections to the ventral striatum (VS)^9,22,39,40^. TS dopamine may interfere indirectly with these processes. There are at least two possible mechanisms through which TS dopamine interacts with appetitive circuitry.

First, TS dopamine and appetitive circuits may converge in brain regions where the balance between avoidance and approach is regulated. Both TS and VS project to the substantia nigra reticulata (SNr), forming the striatonigral pathway, but target distinct subregions; lateral and medial SNr, respectively. Previous studies have shown that these subregions project to different brainstem areas, such as the inferior colliculus and medial superior colliculus^41,42^. Several reports also indicate substantial projections from the SNr to the zona incerta (ZI)^43–45^, which is involved in both aversion-induced avoidance^46–48^ and reward-based approach behaviors^49–52^. In addition to the striatonigral pathway, the striatopallidal pathway—where the TS projects to the globus pallidus externa (GPe) and the VS projects to the ventral pallidum—also relays information to the ZI^53–55^. Investigating whether aversive and appetitive signals converge in the ZI would be valuable.

Second, TS dopamine may influence appetitive circuitry indirectly by modulating anxiety-related brain networks, such as the extended amygdala. Notably, the lateral, but not medial, SNr projects to the basolateral and central amygdala^56^, key regions involved in anxiety regulation^57–59^. Although the GPe has only modest connections to the amygdala^60,61^, the central amygdala and the bed nucleus of the stria terminalis (BNST) are known to communicate reciprocally, a relationship critical for regulating anxiety levels^62,63^. Notably, BNST neurons have been shown to inhibit dopamine neurons involved in conventional reward processing by activating local GABAergic neurons^64–66^, suggesting that anxiety-related circuits may interfere with appetitive learning and behavior.

Using a dual valence-driven classical conditioning task, we examined the roles of TS dopamine in aversive and appetitive learning and behaviors. Our results indicate that reduced TS dopamine signaling is essential for adapting to aversive stimuli and, subsequently, for enhancing appetitive learning. Future studies should further clarify how downstream targets of TS dopamine regulate aversive and appetitive behaviors.

## Methods

### Animals

A total of 35 adult C57BL/6J (Jackson Laboratory) male and female mice were used in these experiments. Nine wild-type animals (eight male and one female) were used for the 2-VCC task. For validation of optogenetic manipulation, eight *DAT-cre* (B6.SJL-Slc6a3^tm1.1^(cre) ^Bkmn^/J, Jackson Laboratory; RRID: IMSR JAX:006660; four male and four female)^67^ heterozygous mice were used. For optogenetic manipulation, we used seven *DAT-cre* (four male and three female) and eleven wild-type mice. Animals were housed on a 12-12 h dark-light cycle and performed the task at the similar timing (±2 h), during the light period. Ambient temperature was kept at 20±3°C, and humidity was kept below 60%. Animals were group-housed (2-6 animals per cage) until surgery, then individually housed throughout training and testing. Some mice were water restricted for behavioral tests. In those cases, mice received water every day by experimenters, and body weights were kept >85% of their original weights. All procedures were approved by the Hokkaido Institutional Animal Care and Use Committee.

### Two-valence classical conditioning task

#### Training

After >2 weeks of recovery from the surgery, water restriction was started and continued until the final day of behavioral testing. The mice were habituated to be head-fixed for 3 days before training. During these days, the mice were head-fixed and given a water drop of either 2, 4, or 8 μl by a water delivery system (OPR-7300, O’ HARA & CO., LTD., Japan) at random inter trial intervals (ITI, exponential distribution between 10 s and 20 s, average 13 s). An odor mask without air delivery was set to their nose on the second day. Same procedure but with air delivery from the odor mask was performed on the third day. Each session consisted of 60 trials.

The mice were then trained with a simple classical conditioning task to learn that an odor (i.e., CS) tells upcoming unconditioned stimulus (i.e., US) ^39,68^. Odors were delivered by an olfactory delivery system (SDC-2060, O’ HARA & CO., LTD., Japan). Each odor was dissolved in mineral oil at 1:10 dilution. 30 μl of diluted odor was applied to a syringe filter (2.7 μm pore, 13 mm diameter; 6823-1327, Whatman). Animals used for behavioral task were conditioned with three different neutral odors, chosen at pseudorandomly from these seven: isoamyl acetate, *p*-cymene, (*R*)-limonene, 1-butanol, 1-hexanol, eugenol, hexanoic acid. Three different odors (odor A-C) were associated with either 100% delivery of water, 40% delivery of water, or no consequence. A trial was initiated by an odor delivery for 1 second followed by water delivery or no consequence with 2 second delay. ITI was applied with exponential distribution. Twenty trials per trial type, total 120 trials per session, one session per day for 5 to 10 days were performed.

#### Test

Another set of 3 different odors (odor D-E) were used for the 2-VCC tests. Each odor was associated with either 100% delivery of 4 μl water, 100% deliver of air-puff stimulation (50 LPM for 100 ms, OPR-AS, O’HARA & CO., LTD., Japan), or no consequence. The ratio of the trial types was set at 3:1:3 for reward trial, aversive trial, and null trial, respectively. Delay time between CS and US was set at 2 second. ITI was applied with exponential distribution. Total 70 trials per session, 1 session per day for 5 consecutive days were performed. The tasks were performed inside a sound attenuate box (MC-050, Muromachi Kikai Co., Ltd., Japan). The mice were allowed to run on a running wheel (S24080, NARISHIGE GROUP, Japan) during the tasks. Behavioral task was controlled by custom software written in LabVIEW (National Instruments, US).

#### Licking behavior recording and analysis

During the tasks, licking was detected by a photoelectric sensor (OPR-LK, O’HARA & CO., LTD., Japan) that produces a change in voltage when the light path is broken. The timing of each lick was detected at the peak of the voltage signal above a threshold. To plot the time course of licking patterns, the lick rate was calculated by a moving average of a 200 ms window.

Licking data were analyzed in custom software written in MATLAB (MathWorks, US) and were shown as mean ± S.E.M., unless otherwise stated. Anticipatory licking was defined as average lick/s during the 2 s delay period between the CS and the US delivery time point.

#### Video capture and analysis

To obtain blinking data, a camera (DMK33UP1300, The Imaging Source, LLC, US) was set to capture a left eye of a head-fixed mouse. We used infrared light (850 nm wavelength, C&M Vision Technologies Inc, US) to illuminate the arena and recorded video at 30 frames per second with H.264 video compression and streaming recording mode. The video was captured using the IC Imaging Control software that accompanies the camera and processed using MATLAB (MathWorks, US). To synchronize the video data and the behavioral data, a 20 ms of brief LED flush, which was delivered every 10 second by the behavioral device, was also captured. The LED was sealed with a black light shielding tape not to deliver light stimulation to the animals.

Blinking data were analyzed custom software written in MATLAB (MathWorks, US) and were shown as mean ± S.E.M., unless otherwise stated. To figure out the eye area of each frame, we binarized the image by setting a certain threshold to separate eye area from the background. We then calculated Δarea/area_0_ (area_0_ = the size of eye area at -2 s to -1 s before each trial start) and used to detect blinking behavior. Anticipatory blinking was defined as average eye area size during the 2 s delay period between the CS and the US delivery time point.

### Surgical procedures

All surgeries were performed under a stereotaxic (custom made by NARISHIGE GROUP, Japan) and aseptic conditions with animals anesthetized with isoflurane (1-2% at 0.5-1.0 l min^-1^, Viatris Healthcare Co., LTD., Japan) which delivered by an inhalation anesthesia equipment (TK-36, Biomachinery, Japan). We used the following coordinates to target injections and/or implantations for TS: bregma -1.7 mm, lateral +3.2 mm, depth -2.4 mm; AS: bregma +0.9 ∼ -0.3 mm, lateral -1.0 or -2.0 mm, depth -2.4 mm; and bilateral SNL: bregma -3.3 mm, lateral ±2.1 mm, depth -3.4 mm (relative to dura). To express GRAB_DA2m_^69^ and tdTomato, we unilaterally injected 200 μl of mixed (3:1) virus solution, AAV9-hsyn-GRAB_DA2m_ (5.0×10^13^ particles per milliliter, YL002009-AV9-PUB, WZ Biosciences Inc., US) and AAV5-CAG-tdTomato (5.0×10^12^ particles per milliliter, #59462-AAV5, addgene, US) into the striatum. To express ChrimsonR in dopamine neurons, we bilaterally injected AAV5-Syn-FLEX-rc [ChrimsonR-tdTomato] (2.1×10^13^ particles per milliliter, #62723-AAV5, addgene, US) ^70^ into the SNL in DAT-cre mice. To conduct a control experiment for optogenetics stimulation, we bilaterally injected AAV5-CAG-tdTomato (5.0×10^12^ particles per milliliter, #59462-AAV5, addgene, US) or AAV8-hSyn-mCherry (2.6×10^13^ particles per milliliter, 114472-AAV8, addgene, US) into the SNL in wild type mice. The glass pipettes were made by using a puller (PC-100, NARISHIGE GROUP, Japan). Virus solution was packed into a glass pipette and injected with 200 nl/min flow rate and the pipette lasted around 10 minutes, after which it was slowly removed to prevent damage to the tissue. We also implanted an optic fiber (200 μm diameter, 3.5 mm, RWD Life Science Co., LTD. China) into the virus injection site. To do this, we first slowly lowered an optic fiber to the striatum or SNL, then attached it to the skull with two types of dental cement (Superbond, 221AABZX00115000, Sun Medical Co., LTD., Japan, Unifast III, 218AABZX00018000, GC Co., Japan). A head chamber (CF-10, NARISHIGE GROUP, Japan) was also implanted to do head-fixed behavioral tasks.

### Photometry recording

#### Photometry recording during 2-VCC task

Photometry signals from fluorescent proteins were recorded with a multi-bundle fiber photometry system (BFPD, Doric Lenses, Canada) which consists of two LED light sources, a LED driver, and two CMOS cameras. An LED driver was used to deliver excitation light from LEDs at 460-490 nm and at 555-570 nm (67-575 μW output at fiber tip). We used an optic fiber (R-FOC-BL200C-39NA, RWD Life Science Co., LTD., Chaina) to stably access deep brain regions and interface with a flexible 5-branching patch cord (BBP5, Doric Lenses, Canada) on the skull. The patch cord simultaneously delivers excitation light and collects dopamine sensor and tdTomato fluorescence emissions. Emissions were detected throughout CMOS camera (500-550 nm, 580-680 nm).

#### Signal analysis

Dopamine sensor (green) and tdTomato (red) signals were analyzed by Danse software (Doric Lenses, Canada). The global slope within a session of was removed using an arPLS ^71^ method. Then, the correlation between green and red signals was examined by linear regression using signals during an entire session head-fixed task. If the correlation was significant (p<0.05), the fitted red signals were subtracted from green signals. Then ΔF/F_0_ was calculated (F_0_ = signals at -2 s to -1 s before each trial start) and standardized as z-score.

To test correlation between dopamine signals and behavior (regression slope of anticipatory blinking or licking over the trials) across individuals or across trials, dopamine responses in each animal were first averaged within a session. The obtained responses in each session in each animal were used to examine time-course of the responses across sessions in each animal. The average responses within a session were further averaged across sessions in each animal to prevent sampling bias across sessions, which was used to examine individual variability. To examine effects of trial types, dopamine responses in aversive trials, reward trials, and null trials in each animal were first averaged within a session, and then averaged across sessions to visualize activity patterns in each trial type. To compare responses between trial types, all trials of each trial type in each animal were averaged and directly compared within animals. The same dataset was also examined with 3-way Analysis of Variance (ANOVA) with animal identity (ID), trial types, and trial number as predictive factors. ID was treated as a random variable and others as fixed, and trial was treated as a continuous variable and others as categorical.

### Optogenetic manipulation

#### Optimization of optogenetic excitation

In order to perform optogenetic manipulation and photometry recording simultaneously, the mice with ChrimsonR or tdTomato in unilateral SNL and GRAB_DA2m_ in ipsilateral TS were used. Red-light was delivered from LED light source (M625F2, Thorlabs, US) to φ200 μm optic fibers (R-FOC-BL200C-39NA, RWD Life Science Co., LTD., China) attached upon the SNL via a φ200 μm bifurcate fiber bundle patch cord (BFYL1LF01, Thorlabs, US). Pulses of 1.4 mW light with a duration of 5 ms were delivered at frequencies of 5, 10, 20, or 40 Hz for 600 ms, controlled by custom software written in LabVIEW (National Instruments, US) via a NIDAQ board (USB-6008, National Instruments, US).

The same equipment and procedures were used for photometry recording (see **Photometry recording**), except for that the tdTomato correction was not applied. Mice were head-fixed and exposed to air puff stimulation to the left eye or the red-light stimulation either 5, 10, 20, or 40 Hz to the SNL. Each trial type consisted of 8 trials. Total 40 trials per session and 3 sessions were performed. Because 40 Hz red light stimulation produced significant artifact to the dopamine signal, further analysis was not performed.

#### Optogenetic excitation of dopamine neurons with photometry recording

The same behavioral training and tests were applied to the mice with ChrimsonR or tdTomato/mCherry in bilateral SNL and GRAB_DA2m_ in unilateral TS (see **Two-valence classical conditioning task**) except for the 10 Hz optogenetic stimulation of SNL dopamine neurons at US_puff_ timing in day 2-5 of 2-VCC task. The parameters and procedures of optogenetic stimulation and photometry recording were exactly same as above.

### Histology and immunohistochemistry

Immunohistochemistry was performed in the similar manner as previously reported^22^. Briefly, animals were perfused using 4% paraformaldehyde and then their brains were sliced into 100 µm thick coronal sections using a vibratome (DTK-1000, DOSAKA EM CO., LTD., Japan) and stored in PBS. To visualize dopamine cell bodies in the midbrain, the brain slices were stained with rabbit anti-tyrosine hydroxylase antibodies (1:500, TH; AB152, MilliporeSigma, US) at 4°C overnight and then with fluorescent secondary antibodies (Alexa Fluor 647, 1:1000, A-21244, Thermo Fisher Scientific, US) at RT for 2 hours. The slices were then mounted in Fluoromount (K024, Diagnostic BioSystems, US) and imaged with a fluorescence microscopy (BZ-X710, KEYENCE CO., Japan) or confocal laser microscopy (LSM900, Carl Zeiss CO., LTD., Germany). In the figures, anatomical positions were plotted on the nearest reference slice ^72,73^.

### Statistics

Data analysis was performed using custom software written in MATLAB (MathWorks, US) and SPSS ver. 26 (IBM, US). All statistical tests were two-sided and had an α level of 0.05. For statistical comparison of mean, we used repeated or 1-way ANOVA, two-sample Student’s t-test, two-sample paired t-test, and one sample t-test, unless otherwise noted. Bonferroni’s method was used for the multiple comparison. All error bars in the figures are S.E.M., unless notification was given. In the boxplots, the edges of the boxes are the 25th and 75th percentiles, and the whiskers extend to the most extreme data points not considered outliers. The major experimental data were taken from more than two separate batch of experiments, and the observed tendency was conserved among the batches.

### Reporting summary

Further information on research design is available in the Nature Portfolio Reporting Summary linked to this article.

## Acknowledgements

We thank all Minami lab members for discussion. We thank Prof. Shinichi Nakagawa at Hokkaido University for sharing his confocal microscopy. This work is supported by Japan Society for the Promotion of Science (JSPS) KAKENHI grant number 25K02415, and PRESTO and JST grant number JPMJPR22S4.

## Author contributions

I.T.-K. and R.T. conceptualized and designed the study. R.T. and I.T.-K. performed the experiments. Y.T. performed animal holding and care. R.T. visualized and analyzed the data. R.T. and I.T.-K wrote the original draft. R.T., Y.T., M.M, and I.T.-K. edited the manuscript. I.T.-K supervised the study.

## Materials & Correspondence

Iku Tsutsui-Kimura

## Data availability

All data, including photometry and the behavioral data, will be available via Dryad after acceptance.

## Supplementary video files

**Video S1. Licking behavior in a 2-VCC task**

Video showing an example of licking behavior in a 2-VCCtask. The mouse licked the water spout after application of USwater.

**Video S2. Blink response to an air-puff stimulation**

Video showing an example of blinking behavior in a 2-VCCtask. The mouse blinked at the timing of air-puff stimulation to the left eye.

**Video S3. Anticipatory blinking in a 2-VCC task**

Video showing an example of anticipatory blinking in a 2-VCCtask. The mouse started to blink after the CSpuff presentation before the USpuff application.

## Reference

1. Goodman, J., McClay, M. & Dunsmoor, J. E. Threat-induced modulation of hippocampal and striatal memory systems during navigation of a virtual environment. Neurobiol. Learn. Mem. 168, 107160 (2020).

2. Silston, B., Ochsner, K. N. & Aly, M. Threat impairs flexible use of a cognitive map. Motiv. Emot. 47, 908–927 (2023).

3. Walters, C. J., Jubran, J., Sheehan, A., Erickson, M. T. & Redish, A. D. Avoid-approach conflict behaviors differentially affected by anxiolytics: implications for a computational model of risky decision-making. Psychopharmacology (Berl.) 236, 2513–2525 (2019).

4. Thornton, I. M., Tagu, J., Zdravković, S. & Kristjánsson, Á. The Predation Game: Does dividing attention affect patterns of human foraging? Cogn. Res. Princ. Implic. 6, 35 (2021).

5. Schultz, W., Dayan, P. & Montague, P. R. A neural substrate of prediction and reward. Science 275, 1593–1599 (1997).

6. Watabe-Uchida, M., Eshel, N. & Uchida, N. Neural Circuitry of Reward Prediction Error. Annu. Rev. Neurosci. 40, 373–394 (2017).

7. Menegas, W. et al. Dopamine neurons projecting to the posterior striatum form an anatomically distinct subclass. eLife 4, e10032 (2015).

8. Akiti, K. et al. Striatal dopamine explains novelty-induced behavioral dynamics and individual variability in threat prediction. Neuron 110, 3789–3804.e9 (2022).

9. Menegas, W., Babayan, B. M., Uchida, N. & Watabe-Uchida, M. Opposite initialization to novel cues in dopamine signaling in ventral and posterior striatum in mice. eLife 6, e21886 (2017).

10. Menegas, W., Akiti, K., Amo, R., Uchida, N. & Watabe-Uchida, M. Dopamine neurons projecting to the posterior striatum reinforce avoidance of threatening stimuli. Nat. Neurosci. 21, 1421–1430 (2018).

11. Tsutsui-Kimura, I. et al. Dopamine in the tail of the striatum facilitates avoidance in threat–reward conflicts. Nat. Neurosci. 28, 795–810 (2025).

12. Zafiri, D., Salinas-Hernández, X. I., De Biasi, E. S., Rebelo, L. & Duvarci, S. Dopamine prediction error signaling in a unique nigrostriatal circuit is critical for associative fear learning. Nat. Commun. 16, 3066 (2025).

13. Kim, H. F., Ghazizadeh, A. & Hikosaka, O. Dopamine Neurons Encoding Long-Term Memory of Object Value for Habitual Behavior. Cell 163, 1165–1175 (2015).

14. Hikosaka, O., Ghazizadeh, A., Griggs, W. & Amita, H. Parallel basal ganglia circuits for decision making. J. Neural Transm. 125, 515–529 (2018).

15. Chen, A. P. F. et al. Nigrostriatal dopamine pathway regulates auditory discrimination behavior. Nat. Commun. 13, 5942 (2022).

16. Matsumoto, H., Tian, J., Uchida, N. & Watabe-Uchida, M. Midbrain dopamine neurons signal aversion in a reward-context-dependent manner. eLife 5, e17328 (2016).

17. Dai, J. & Sun, Q.-Q. Distinct Roles of Somatostatin and Parvalbumin Interneurons in Regulating Predictive Actions and Emotional Responses During Trace Eyeblink Conditioning. 2025.03.23.644831 Preprint at 10.1101/2025.03.23.644831 (2025).

18. Amo, R. et al. A gradual temporal shift of dopamine responses mirrors the progression of temporal difference error in machine learning. Nat. Neurosci. 25, 1082– 1092 (2022).

19. Glimcher, P. W. Understanding dopamine and reinforcement learning: The dopamine reward prediction error hypothesis. Proc. Natl. Acad. Sci. 108, 15647–15654 (2011).

20. Schultz, W. Updating dopamine reward signals. Curr. Opin. Neurobiol. 23, 229– 238 (2013).

21. Schultz, W. Dopamine reward prediction error coding. Dialogues Clin. Neurosci. 18, 23–32 (2016).

22. Tsutsui-Kimura, I. et al. Dopamine in the tail of the striatum facilitates avoidance in threat–reward conflicts. Nat. Neurosci. 28, 795–810 (2025).

23. Chen, A. P. F. et al. Nigrostriatal dopamine modulates the striatal-amygdala pathway in auditory fear conditioning. Nat. Commun. 14, 7231 (2023).

24. Maren, S. & Quirk, G. J. Neuronal signalling of fear memory. Nat. Rev. Neurosci. 5, 844–852 (2004).

25. Myers, K. M. & Davis, M. Behavioral and Neural Analysis of Extinction. Neuron 36, 567–584 (2002).

26. Quirk, G. J. & Mueller, D. Neural Mechanisms of Extinction Learning and Retrieval. Neuropsychopharmacology 33, 56–72 (2008).

27. Rescorla, R. A. Evidence for ‘unique stimulus’ account of configural conditioning. J. Comp. Physiol. Psychol. 85, 331–338 (1973).

28. Solomon, R. L. & Corbit, J. D. An opponent-process theory of motivation: I. Temporal dynamics of affect. Psychol. Rev. 81, 119–145 (1974).

29. Farassat, N. et al. In vivo functional diversity of midbrain dopamine neurons within identified axonal projections. eLife 8, e48408 (2019).

30. Evans, R. C., Zhu, M. & Khaliq, Z. M. Dopamine Inhibition Differentially Controls Excitability of Substantia Nigra Dopamine Neuron Subpopulations through T-Type Calcium Channels. J. Neurosci. 37, 3704–3720 (2017).

31. Lerner, T. N. et al. Intact-Brain Analyses Reveal Distinct Information Carried by SNc Dopamine Subcircuits. Cell 162, 635–647 (2015).

32. Ilango, A., Kesner, A. J., Broker, C. J., Wang, D. V. & Ikemoto, S. Phasic excitation of ventral tegmental dopamine neurons potentiates the initiation of conditioned approach behavior: parametric and reinforcement-schedule analyses. Front. Behav. Neurosci. 8, (2014).

33. Saunders, B. T., Richard, J. M., Margolis, E. B. & Janak, P. H. Dopamine neurons create Pavlovian conditioned stimuli with circuit-defined motivational properties. Nat. Neurosci. 21, 1072–1083 (2018).

34. Rossi, M. A., Sukharnikova, T., Hayrapetyan, V. Y., Yang, L. & Yin, H. H. Operant Self-Stimulation of Dopamine Neurons in the Substantia Nigra. PLOS ONE 8, e65799 (2013).

35. Keiflin, R., Pribut, H. J., Shah, N. B. & Janak, P. H. Ventral Tegmental Dopamine Neurons Participate in Reward Identity Predictions. Curr. Biol. 29, 93–103.e3 (2019).

36. Zigmond, M. J., Hastings, T. G. & Perez, R. G. Increased dopamine turnover after partial loss of dopaminergic neurons: compensation or toxicity? Parkinsonism Relat. Disord. 8, 389–393 (2002).

37. Coddington, L. T., Lindo, S. E. & Dudman, J. T. Mesolimbic dopamine adapts the rate of learning from action. Nature 614, 294–302 (2023).

38. Long, C. et al. Constraints on the subsecond modulation of striatal dynamics by physiological dopamine signaling. Nat. Neurosci. 27, 1977–1986 (2024).

39. Amo, R., Uchida, N. & Watabe-Uchida, M. Glutamate inputs send prediction error of reward but not negative value of aversive stimuli to dopamine neurons. Neuron 112, 1001–1019.e6 (2024).

40. Kim, H. F., Griggs, W. S. & Hikosaka, O. Long-Term Value Memory in the Primate Posterior Thalamus for Fast Automatic Action. Curr. Biol. 30, 2901–2911.e3 (2020).

41. Foster, N. N. et al. The mouse cortico–basal ganglia–thalamic network. Nature 598, 188–194 (2021).

42. McElvain, L. E. et al. Specific populations of basal ganglia output neurons target distinct brain stem areas while collateralizing throughout the diencephalon. Neuron 109, 1721–1738.e4 (2021).

43. Shammah-Lagnado, S. J., Negrão, N. & Ricardo, J. A. Afferent connections of the zona incerta: A horseradish peroxidase study in the rat. Neuroscience 15, 109–134 (1985).

44. Kolmac, C. I., Power, B. D. & Mitrofanis, J. Patterns of connections between zona incerta and brainstem in rats. J. Comp. Neurol. 396, 544–555 (1998).

45. Heise, C. E. & Mitrofanis, J. Evidence for a glutamatergic projection from the zona incerta to the basal ganglia of rats. J. Comp. Neurol. 468, 482–495 (2004).

46. Wang, X. et al. A cross-modality enhancement of defensive flight via parvalbumin neurons in zona incerta. eLife 8, e42728 (2019).

47. Chou, X. et al. Inhibitory gain modulation of defense behaviors by zona incerta. Nat. Commun. 9, 1151 (2018).

48. Zhou, H., Xiang, W. & Huang, M. Inactivation of Zona Incerta Blocks Social Conditioned Place Aversion and Modulates Post-traumatic Stress Disorder-Like Behaviors in Mice. Front. Behav. Neurosci. 15, (2021).

49. Zhang, X. & van den Pol, A. N. Rapid binge-like eating and body weight gain driven by zona incerta GABA neuron activation. Science 356, 853–859 (2017).

50. Zhao, Z. et al. Zona incerta GABAergic neurons integrate prey-related sensory signals and induce an appetitive drive to promote hunting. Nat. Neurosci. 22, 921–932 (2019).

51. Kendrick, K. M. & Baldwin, B. A. Characterization of neuronal responses in the zona incerta of the subthalamic region of the sheep during ingestion of food and liquid. Neurosci. Lett. 63, 237–242 (1986).

52. Kendrick, K. M. & Baldwin, B. A. The effects of sodium appetite on the responses of cells in the zona incerta to the sight or ingestion of food, salt and water in sheep. Brain Res. 492, 211–218 (1989).

53. Gorbachevskaya, A. I. & Chivileva, O. G. Structural organization of the zona incerta of the dog diencephalon. Neurosci. Behav. Physiol. 38, 573–8 (2008).

54. Gorbachevskaya, A. I. Connections between the Zona Incerta of the Dog Diencephalon and the Substantia Nigra, Ventral Tegmental Field, and Pedunculopontine Tegmental Nucleus. Neurosci. Behav. Physiol. 40, 603–7 (2010).

55. Arena, G. et al. Disentangling the identity of the zona incerta: a review of the known connections and latest implications. Ageing Res. Rev. 93, 102140 (2024).

56. Beckstead, R. M., Domesick, V. B. & Nauta, W. J. H. Efferent connections of the substantia nigra and ventral tegmental area in the rat. Brain Res. 175, 191–217 (1979).

57. Sengupta, A. et al. Basolateral Amygdala Neurons Maintain Aversive Emotional Salience. J. Neurosci. 38, 3001–3012 (2018).

58. LeDuke, D. O., Borio, M., Miranda, R. & Tye, K. M. Anxiety and depression: A top-down, bottom-up model of circuit function. Ann. N. Y. Acad. Sci. 1525, 70–87 (2023).

59. Ludkiewicz, B., Pszczolinska, A., Moryś, J. & Kowiański, P. The rodent amygdala under acute psychological stress: a review. Folia Morphol. 0, (2025).

60. Courtney, C. D., Pamukcu, A. & Chan, C. S. Cell and circuit complexity of the external globus pallidus. Nat. Neurosci. 26, 1147–1159 (2023).

61. Giossi, C., Rubin, J. E., Gittis, A., Verstynen, T. & Vich, C. Rethinking the external globus pallidus and information flow in cortico-basal ganglia-thalamic circuits. Eur. J. Neurosci. 60, 6129–6144 (2024).

62. Asok, A. et al. Optogenetic silencing of a corticotropin-releasing factor pathway from the central amygdala to the bed nucleus of the stria terminalis disrupts sustained fear. Mol. Psychiatry 23, 914–922 (2018).

63. Gungor, N. Z. & Paré, D. Functional Heterogeneity in the Bed Nucleus of the Stria TerminalisFunctional Heterogeneity in the Bed Nucleus of the Stria Terminalis. J. Neurosci. 36, 8038–8049 (2016).

64. Jennings, J. H. et al. Distinct extended amygdala circuits for divergent motivational states. Nature 496, 224–228 (2013).

65. Kudo, T. et al. GABAergic neurons in the ventral tegmental area receive dual GABA/enkephalin-mediated inhibitory inputs from the bed nucleus of the stria terminalis. Eur. J. Neurosci. 39, 1796–1809 (2014).

66. Minami, M. & Ide, S. How Does Pain Induce Negative Emotion? Role of the Bed Nucleus of the Stria Terminalis in Pain-Induced Place Aversion. Curr. Mol. Med. 15, 184– 190 (2015).

67. Bäckman, C. M. et al. Characterization of a mouse strain expressing Cre recombinase from the 3’ untranslated region of the dopamine transporter locus. Genes. N. Y. N 2000 44, 383–390 (2006).

68. Cai, X. et al. Dopamine dynamics are dispensable for movement but promote reward responses. Nature 635, 406–414 (2024).

69. Sun, F. et al. Next-generation GRAB sensors for monitoring dopaminergic activity in vivo. Nat. Methods 17, 1156–1166 (2020).

70. Klapoetke, N. C. et al. Independent optical excitation of distinct neural populations. Nat. Methods 11, 338–346 (2014).

71. Baseline correction using asymmetrically reweighted penalized least squares smoothing - Analyst (RSC Publishing) DOI:10.1039/C4AN01061B. https://pubs.rsc.org/en/content/articlehtml/2014/an/c4an01061b.

72. Keith, B. J., Franklin, G. P. & Paxinos, G. The mouse brain in stereotaxic coordinates. Calif. Acad. (2008).

73. Franklin, K. B. J. & Paxinos, G. Paxinos and Franklin’s the Mouse Brain in Stereotaxic Coordinates, Compact: The Coronal Plates and Diagrams. (Academic Press, 2019).

