## Supplemental Figures for "Competing valence-related roles of dopamine in the tail of the striatum"

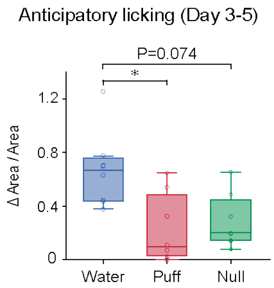


**Figure S1. Increased anticipator licking in the later sessions**

Mice discriminatively showed anticipatory licking in the later sessions of appetitive trials ($p=3.0\times10^{-3}$, $F(2, 14)=8.8$, repeated ANOVA; $p=0.045$, aversive vs appetitive, $p=0.82$, aversive vs null, $p=0.074$, appetitive vs null, Bonferroni's test, $n=8$ animals for each).

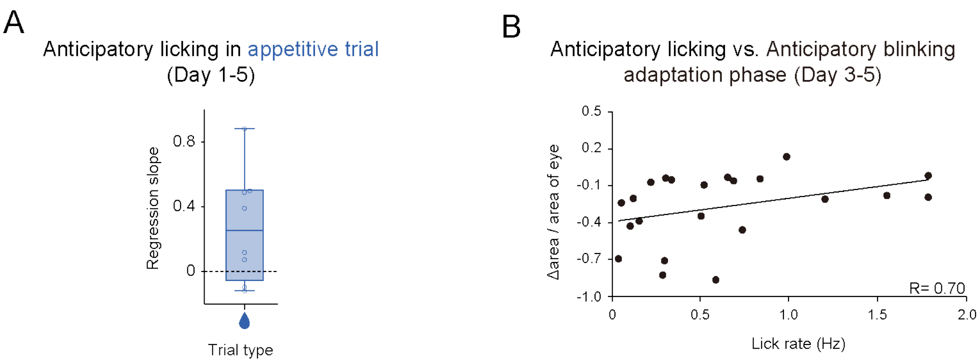


**Figure S2. Two competing valence-induced learning in the 2-VCC task**

**A**, No significant trend in regression coefficients of the rate of anticipatory licking with trial number in Day 1-5 for each animal (p=0.056, one-sampled t-test, n=8). **B**, Regression coefficients of intensities of anticipatory blinking with anticipatory licking were correlated in the adaptation phase (R=0.70, p=0.026, Pearson's correlation coefficient, n=22). Error bars, SEM. Center of box plot shows median; edges are 25th and 75th percentiles; and whiskers are the most extreme data points. *p < 0.05.

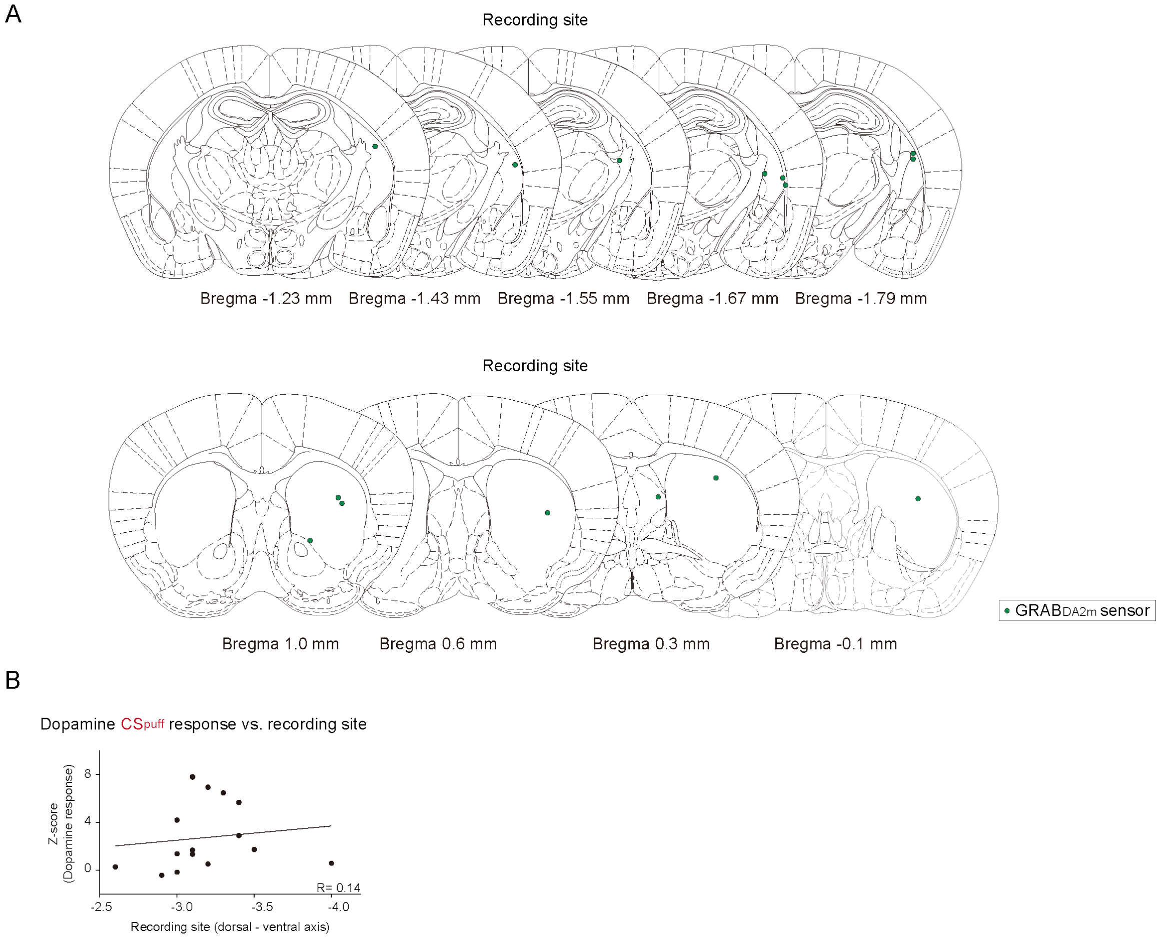


### Figure S3. Striatal dopamine dynamics during the 2-VCC task

**A**, Location of optic fiber tips (marked with green circles) used to collect dopamine sensor signals in the striatum. **B**, Dopamine CSpuff response was not correlated with dorso-ventral axis of recording location (R=0.14, p=0.63, Pearson's correlation coefficients, n=15 animals).

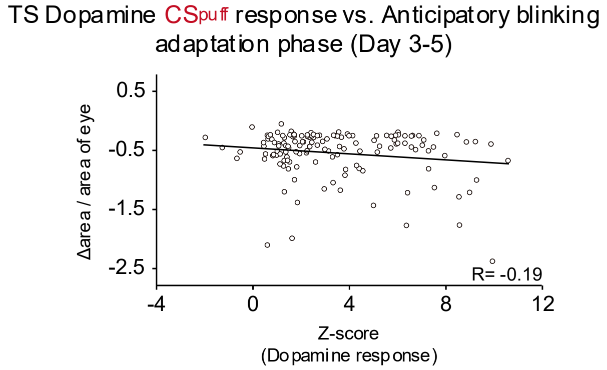


**Figure S4. Correlation between CSpuff response and anticipatory blinking**

Regression coefficients of intensities of anticipatory blinking with amplitude of TS dopamine CSpuff response were anti-correlated (R=-0.19, p=0.025, Pearson's correlation coefficient, n=140).

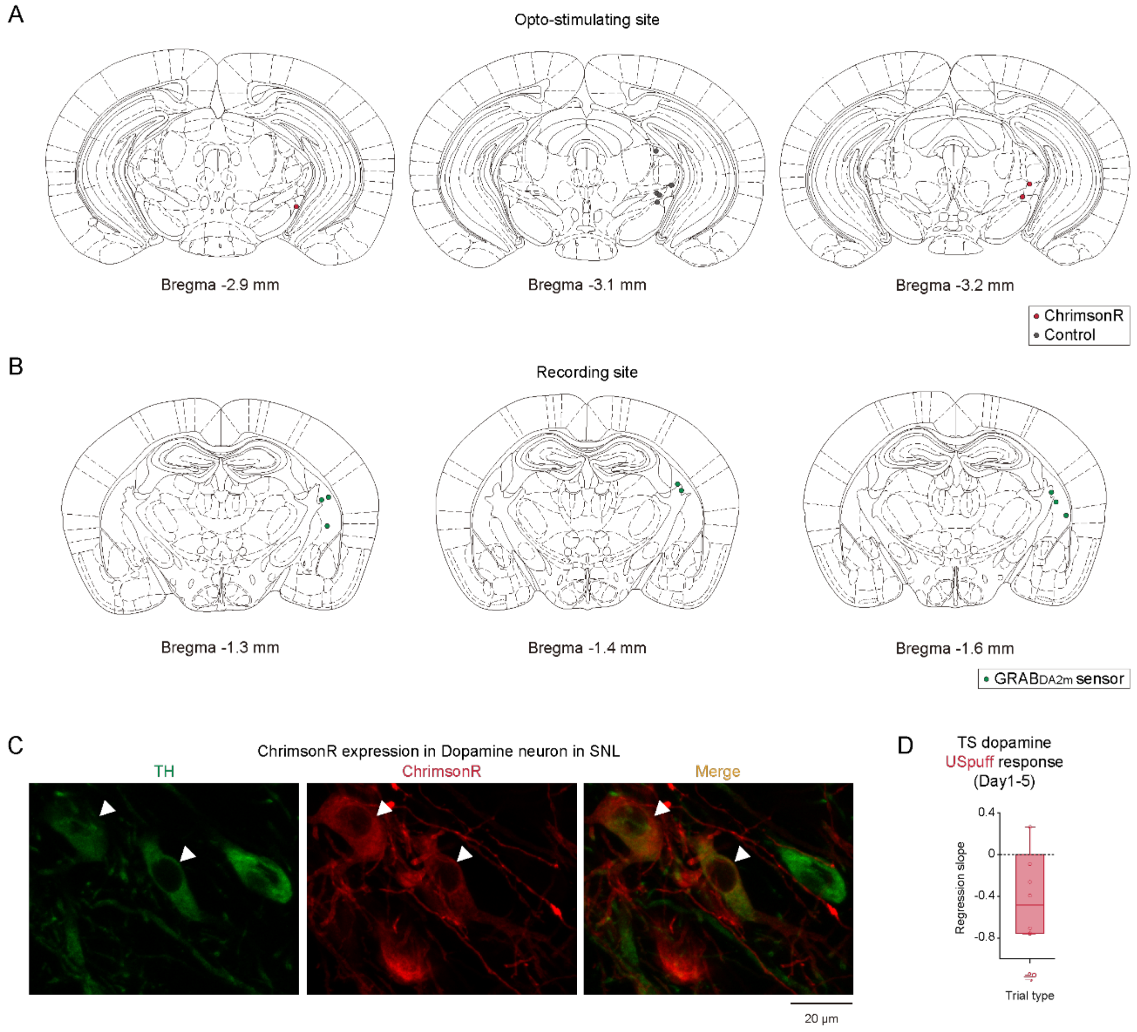


### Figure S5. Validation for optogenetic manipulation of SNL dopamine neurons

**A,** Location of optic fiber tips (ChrimsonR group was marked with red circles while control group was marked with gray circles) used to opto-stimulate ChrimsonR in the SNL. **B,** Location of optic fiber tips (marked with green circles) used to collect dopamine sensor signals in the striatum. **C,** TH (green) and ChrimsonR (red) expression in DAT-cre mouse. White arrow heads indicate cells that express both. Scale bar, 20 μm. **D**, Regression coefficients of USpuff dopamine response with trial number in Day 1-5 for each animal were not significantly changed (p=0.077, one-sampled t-test, n=6). Error bars, SEM. Center of box plot shows median; edges are 25th and 75th percentiles; and whiskers are the most extreme data points. *p < 0.05.

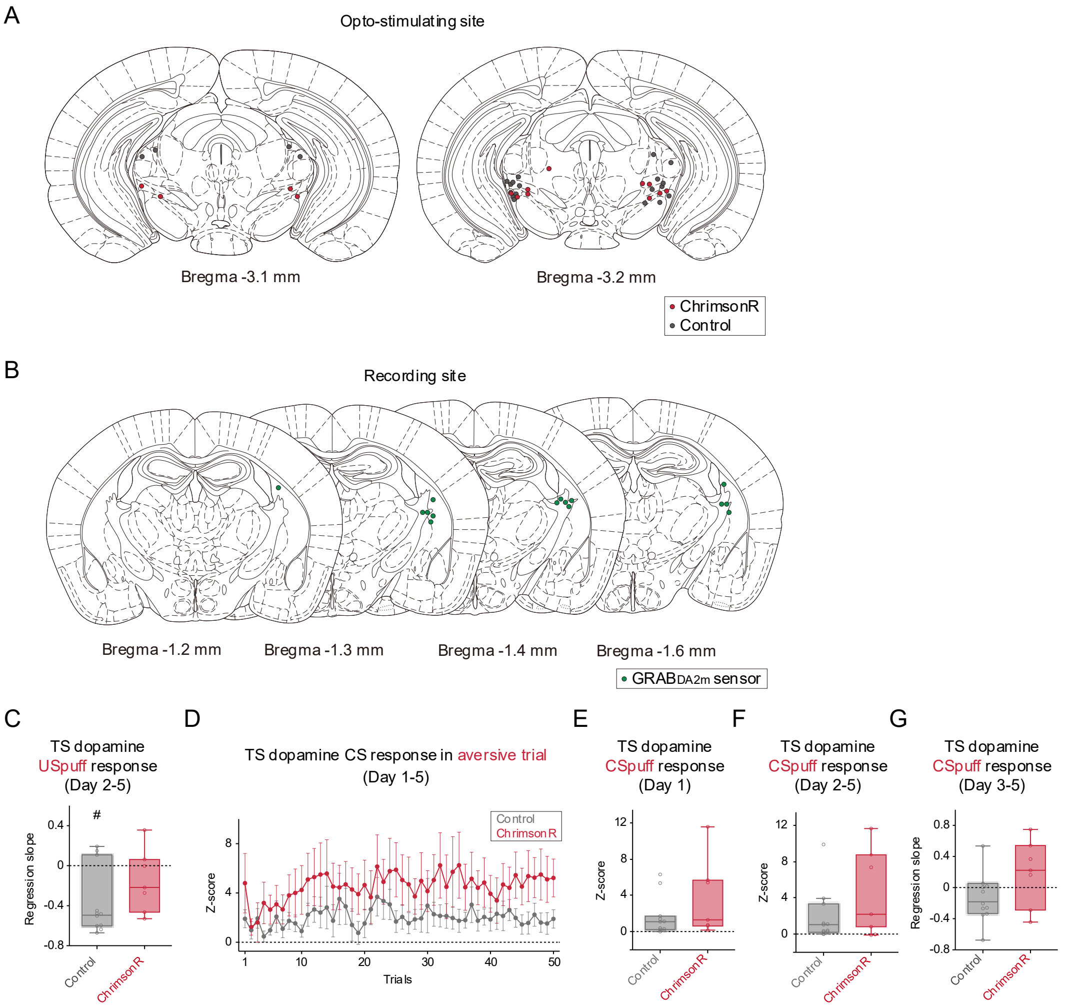


### Figure S6. Optogenetic manipulation affected aversive and appetitive learning

**A**, Location of optic fiber tips (ChrimsonR group was marked with red circles while control group was marked with gray circles) used to opto-stimulate ChrimsonR in the SNL. **B,** Location of optic fiber tips (marked with green circles) used to collect dopamine sensor signals in the striatum. **C**, Regression coefficients of USpuff dopamine response with trial number in Day 2-5 for each animal were significantly negative in the control ($p=5.1\times10^{-3}$, one-sampled t-test, n=11) but not in ChrimsonR group (p=0.24, one-sampled t-test, n=7). **D**, Time-course of CSpuff dopamine response in the aversive trials. **E**, No difference in CSpuff response in control and ChrimsonR groups in Day 1 (p=0.21, two-sided t-test, control group, n=11, ChrimsonR group, n=7). **F**, CSpuff response was not changed by optogenetic stimulation (p=0.18, two-sided t-test, control group, n=11, ChrimsonR group, n=7). **G**, Regression coefficients of CSpuff dopamine response with trial number in Day 3-5 for each animal were relatively high in the ChrimsonR group ($p=0.089$, two-sided t-test, control group, $n=11$, ChrimsonR group, $n=7$).
